# Catechol-Containing Compounds are a Broad Class of Amyloid Inhibitors: Redox State is a Key Determinant of the Inhibitory Activities

**DOI:** 10.1101/2020.10.08.873620

**Authors:** Paul Velander, Ling Wu, Sherry B. Hildreth, Nancy J. Vogelaar, Biswarup Mukhopadhyay, Shijun Zhang, Richard F. Helm, Bin Xu

**Author notes:** Equal contribution (Paul Velander, Ling Wu).

## Abstract

Mechanisms of amyloid inhibition remains poorly understood, in part because most protein targets of amyloid assembly are partially unfolded or intrinsically disordered, which hinders detailed structural characterization of protein-inhibitor complexes and structural-based mechanistic elucidation. Herein we employed a small molecule screening approach to identify inhibitors against three prototype amyloidogenic proteins: amylin, Aβ and tau. One remarkable class of inhibitors identified was catechol-containing compounds and redox-related quinones/anthraquinones. Further mechanistic studies determined that the redox state of the broad class of catechol-containing compounds is a key determinant of the amyloid inhibitor activities.

---

Amyloid diseases represent some of the most debilitating and increasingly more common aging-related human diseases. Alzheimer’s, Parkinson’s, prion diseases, and type 2 diabetes all fall under this category (Westermark et al, 2011; Knowles et al, 2014; Chiti & Dobson, 2017; Wang et al, 2019; Wu et al, 2019). While there are currently no FDA approved anti-amyloid drugs available, data over the years continue to support the notion that one of the most attractive therapeutic targets against amyloidosis are the misfolding amyloid proteins themselves (Eisenberg & Jucker, 2012; Selkoe & Hardy, 2016; Li & Gotz, 2017). Insights gained from several latest high-resolution amyloid structures, and atomic details on their interactions with amyloid inhibitors have provided potentially exciting new paths towards the identification, characterization, and optimization of clinically viable inhibitor candidates (Fitzpatrick et al, 2017; Krotee et al, 2017; Li et al, 2017; Seidler et al, 2018; Spanopoulou et al, 2018; Griner et al, 2019; Armiento et al, 2020). Key to this development is achieving a better understanding of amyloid inhibition mechanisms and processes. Recent investigations have begun to elucidate some of the covalent and non-covalent mechanisms for small molecules or peptide-based inhibitors interacting with amyloidogenic proteins at various stages of amyloid formation (Velander et al, 2017; Seidler et al, 2018; Armiento et al, 2019; Griner et al, 2019). Examples include stabilizing the soluble native confirmations of some amyloidogenic proteins (Johnson et al, 2012), perturbing or “remodeling” unaggregated or pre-aggregated amyloid species towards forming presumably innocuous, non-amyloidogenic aggregates (Ehrnhoefer et al, 2008; Hong et al, 2008; Bieschke et al, 2010), or even accelerating amyloid formation (Jha et al, 2016). However, in many cases, chemical classes of the inhibitors and a clear link between the chemical mode of action of an inhibitor and its macromolecular partner are poorly defined. Moreover, the biochemical, biophysical and pharmacological underpinnings of inhibitor-perturbed amyloid aggregation pathways or alternative aggregate assemblies are not always clear.

To systematically identify broad classes of small molecules and chemical types of amyloid inhibitors, we screened a US National Institutes of Health Clinical Collection (NIHCC) drug-repurposing library of 700 drugs and/or investigational compounds with diverse molecular scaffolds or chemical structures. We employed the thioflavin T (ThT) fluorescence-based assay to monitor toxic amyloid formation utilizing a 384-well plate semi high throughput screening platform (Cao et al, 2012; Brumshtein et al, 2015; Seidler et al, 2018; Fig. S1D). To make our findings broadly applicable, we screened in parallel against three prototype amyloidogenic proteins: amylin, Aβ (Figure S1A) as well as 2N4R tau (Spillantini & Goedert, 2013). Amylin amyloidosis and plaque deposition in the pancreas are hallmark features of type 2 diabetes (Verchere et al, 1996; Westermark et al, 2011), whereas Aβ plaques and tau neurofibrillary tangles are well-established neuropathogenic biomarkers for Alzheimer’s disease.

One of the salient findings from the screening is that catechols and redox-related quinones/anthraquinones represent a broad class of amyloid inhibitors. As shown in Fig. 1A, these molecules made up a substantial portion of the identified strong inhibitors, which were defined as exhibiting greater than three standard deviation units below the ThT RFU observed for buffer treated individual amyloidogenic protein controls (dotted line). Fully 13 out of 41 strong inhibitors (32%) for amylin, 11 out of 22 (50%) for tau (2N4R isoform), and 14 out of 29 (48%) for Aβ were catechols or quinone/anthraquinones (highlighted as red dots). In the screens against amylin amyloid, out of 22 catechols and quinones/anthraquinones from the NIHCC library, 21 of them exhibited significant amyloid inhibitory activities (Table S1) with isoproterenol being the exception, with only a weak inhibitory effect. The majority of these catechols and quinones/anthraquinones (16 out of 22 drugs) were further validated by an orthogonal biochemical assay, Photo-Induced Cross-linking of Unmodified Proteins (PICUP), which identified cross-linked oligomers (Figs. S1B and S3A), or by a biophysical method, transmission electron microscopy (TEM; Fig. 1C, Fig. S1C and Table S1).

**Figure 1.**
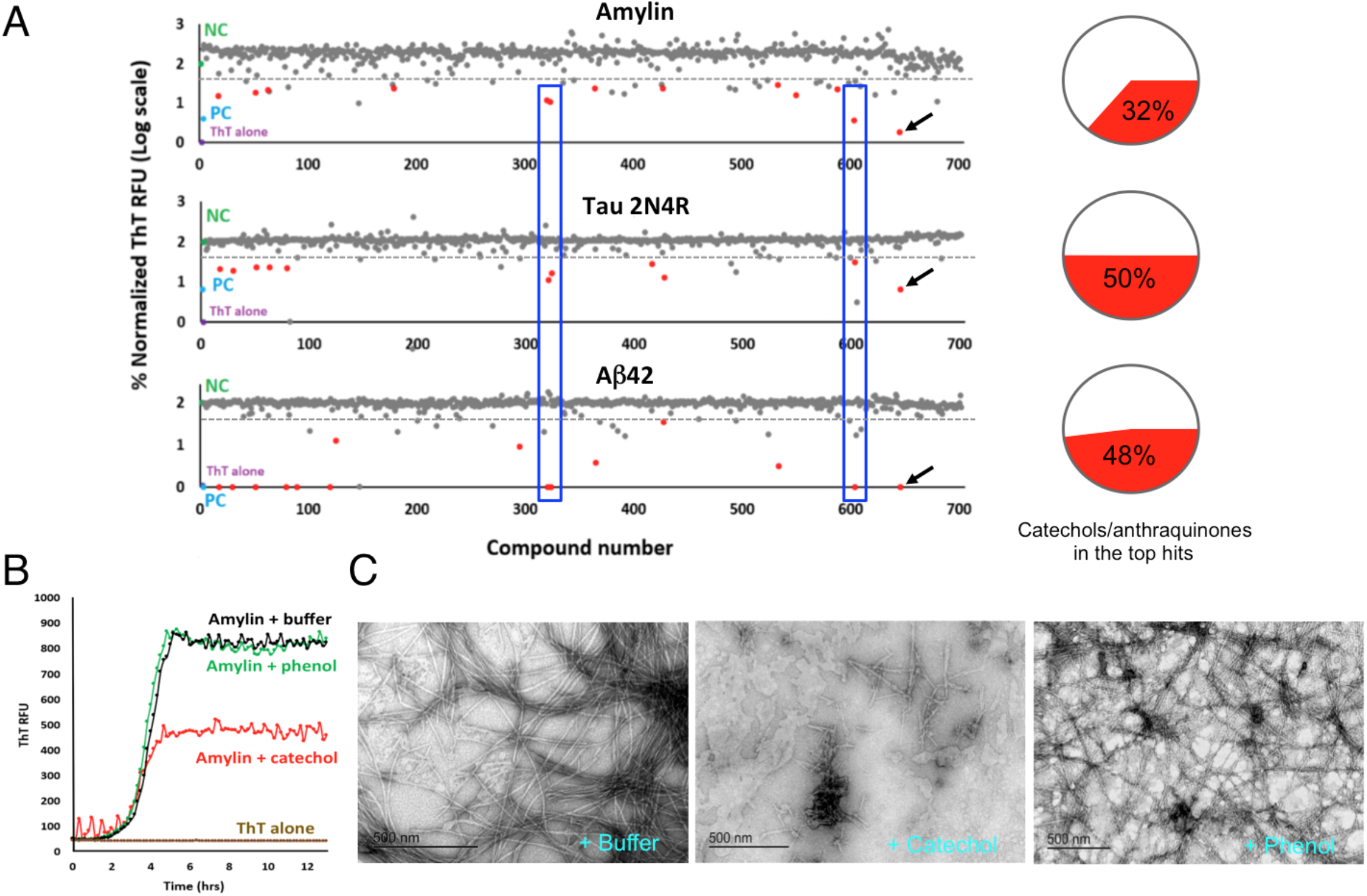
Diverse drug library screening identifies catechols and redox-related anthraquinones as a broad class of amyloid inhibitors. (A) Catechol-containing compounds or redox-related anthraquinones (shown as red dots) were observed to comprise 32% of the top hits (13 out of total 41 top hits) against amylin amyloid formation, 50% of the top hits (11 out of total 22 top hits) against 2N4R isoform of human tau protein, and 48% of the top hits (14 out of total 29 top hits) against Aβ42 amyloid. Top hits were defined by exhibiting a ThT fluorescent signal ≥ three standard deviations below buffer treated negative controls. Each dot represents a drug in the NIH Clinical Collection drug library (total 700 compounds). Buffer treated negative controls (NC), are shown in green. Positive controls (PC), amyloidogenic proteins treated with EGCG, are shown in blue. ThT alone, the assay background, is shown in cyan. Notice y-axis is shown in log-scale. A drug with a value of 2 in y-axis in the figure indicates that it has no inhibitory activity. Examples of common hits (drugs #318, #321, and #601) for all three amyloid proteins are shown in blue boxes. One compound in the library was EGCG (red dots with black arrows), providing collaborating evidence of consistency of the screening method. (B) Catechol, but not phenol, demonstrates moderate yet significant anti-amylin amyloid effects as exhibited by ThT fluorescence assay. (C) TEM analyses validate that catechol, but not phenol, significantly reduces amylin fibril formation. Catechol’s effect is moderate as reflected by the presence of some short fibrils. Typical molar ratios of compound to amyloidogenic protein were 2:1 for all ThT assays and 50:1 for the TEM assays.

Similar with amylin screening results, 17 out of 22 catechols and quinones/anthraquinones showed significant activities against tau 2N4R amyloid (Table S2), with a majority of them further validated by a ThT fluorescence-based tau amyloid remodeling assay, accomplished by spiking testing compounds into pre-formed tau amyloids (Fig. S3B). Fully 18 out of 22 catechols and quinones/anthraquinones exhibited significant inhibition against Aβ amyloid, with nine of them validated previously by assays including TEM and atomic force microscopy (AFM) (Table S3). Chemical structures of catechols and quinones/anthraquinones of the top hits are shown (Fig. S2). Numerous hits such as idarubicin (Compound #318), daunorubicin (Compound #321), and rifapentine (Compound #601), showed strong inhibitory effects to all three amyloidogenic proteins (boxed red dots in Fig. 1A), whereas a few hits displayed preferential inhibition, such as rutin (Compound #548; Tables S1–S3), which preferentially inhibited amylin amyloid formation.

To test the hypothesis whether the catechol functional group alone inhibits amyloid formation, we tested catechol in secondary assays and compared it with its control analog phenol. Catechol demonstrated moderate, yet significant activities in amylin amyloid inhibition in both ThT fluorescence assays and TEM analyses, whereas phenol showed no such inhibitory effects in both assays (Figs. 1B and 1C). These data demonstrate that the catechol functional moiety possesses general anti-amyloid activities. Our finding is supported not only by individual catechol-containing inhibitor examples (Caruana et al, 2011; Sato et al, 2013; Velander et al, 2016; Wu et al, 2017), but also by a large-scale computational data mining study: In a fragment-based combinatorial library screening to identify molecular scaffolds to target certain amyloidogenic proteins in neurodegeneration (Joshi et al, 2016), the catechol group was identified as the fragment with the highest observed occurrence, present in nearly 4,500 compounds among the 16,850 (27%) in the Aβ small molecule library.

Another class of chemical structures enriched in our screens was the anthraquinones/quinones (redox-related to catechols) and tetracyclines. Anthraquinones were previously observed to inhibit tau aggregation (Pickhardt et al, 2005) and quinones were reported to inhibit insulin oligomerization as well as fibril formation (Gong et al, 2014). With respect to tetracycline, our screen revealed that several variants were active in amyloid inhibition. This class of compounds was reported to also inhibit Aβ and β_2_-microglobulin amyloid fibrils (Forioni et al, 2001; Giorgetti et al, 2010).

Based on the fact that catechol-containing compounds and multiple anthraquinone/quinone compounds (redox related to catechols) exhibited strong anti-amyloid activities, we hypothesized that catechol autoxidation may be part of the general mechanism that significantly enhances the anti-amyloid activities of the catechol-containing compounds. To test this hypothesis, we compared the anti-amyloid activities of a collection of oxidized (or aged - exposed to the air for 48 hours) with non-oxidized (non-aged or freshly prepared) catechol-containing compounds. In virtually all cases, aged samples exhibited significantly greater activities than their identically prepared non-aged counterparts (Fig. 2A). Oxidation-induced activity enhancement was not observed with phenol nor a structurally similar but non-catechol amyloid inhibitor, morin. These combined results strongly suggested a catechol-dependent enhancement specificity, which was recapitulated under stringently defined aerobic/anaerobic conditions using an anaerobic chamber. Aerobic, but not anaerobic conditions, significantly enhanced anti-amyloid activities of several catecholamines and other catechol-containing inhibitors (Fig. 2D). The kinetic profiles of ThT fluorescence-based amylin amyloid inhibition showed significantly stronger inhibition with aged RA versus non-aged RA, with an even more dramatic inhibition activity enhancement was observed with aged norepinephrine (Figs. 2B & 2C). Consistently, enhanced inhibition by norepinephrine occurred only under aerobic conditions (Fig. 2E), with marked reduction in fibril formation (Fig. 2F).

**Figure 2.**
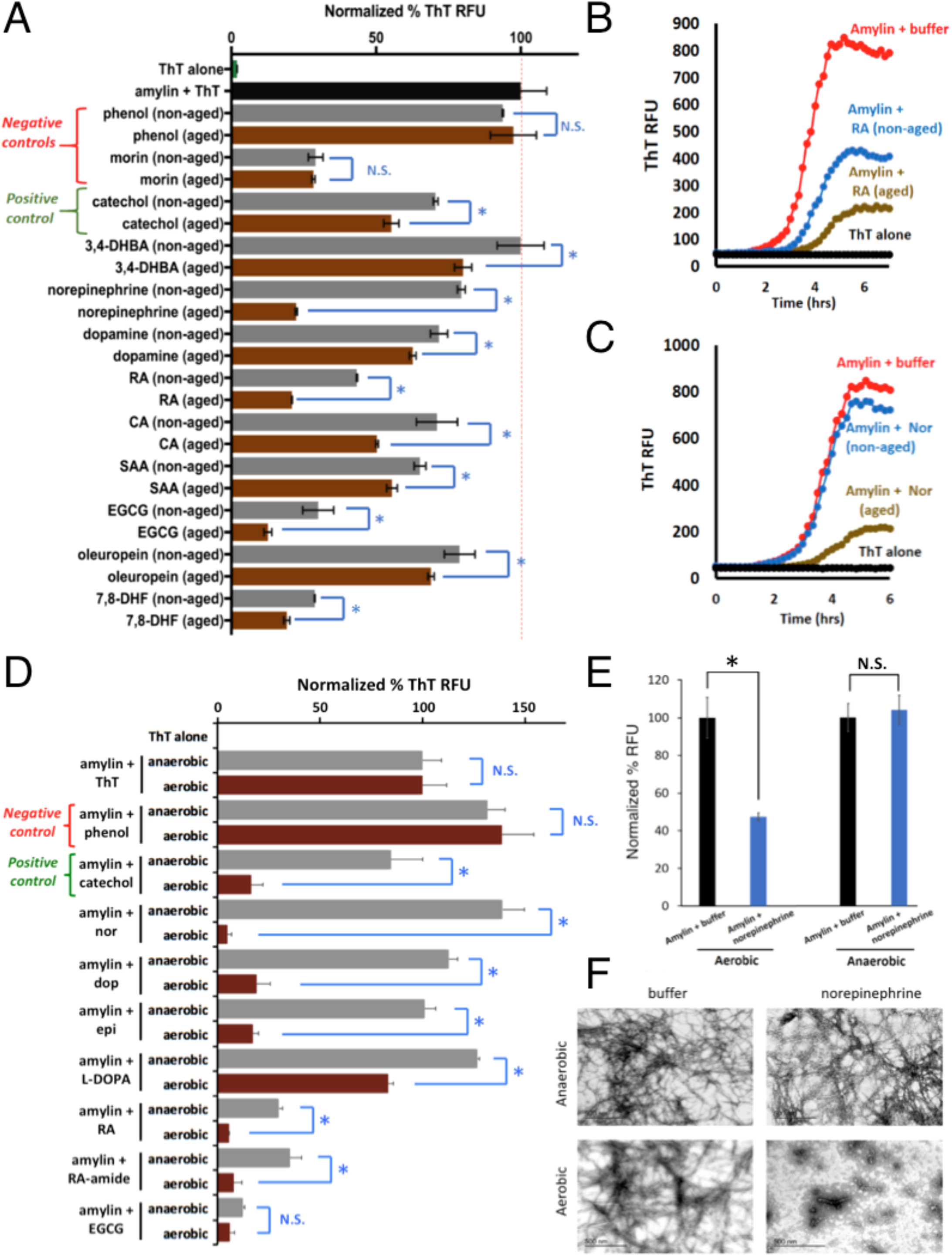
Autoxidation (or aerobic exposure), but not anaerobic conditions, significantly enhanced anti-amyloid activities associated with catechol-containing compounds. (A) Aged catechol-containing compounds (in brown), but not negative controls phenol (a non-inhibitor) or morin (an amylin amyloid inhibitor but with no catechol moiety) display enhanced anti-amyloid activities compared to their non-aged counterparts (in grey). All aged and non-aged treated amylin aggregation reactions were normalized to buffer treated amylin controls (black bar) indicated by the dashed red line. Abbreviations: 7,8-dihydroxy flavone (7,8-DHF), rosmarinic acid (RA), caffeic acid (CA), salvanic acid A (SAA), 3,4-dihydroxybenzoic acid (3,4-DHBA). Statistics were conducted using one-way ANOVA, multiple comparisons test; *p <0.05. (B,C) Representative ThT fluorescence assay time course of aged versus non-aged RA (panel B) and norepinephrine (panel C) treatments in amylin aggregations reactions shown in panel A. (D) Catechol-containing compounds (in brown), showed significantly higher anti-amyloid activities under aerobic exposure compared to their anaerobic condition treated counterparts (in grey). All aerobic exposed and anaerobic condition treated amylin aggregation reactions were normalized to buffer treated amylin controls (amylin + ThT). Phenol served as a negative control and catechol as a positive control. Abbreviations: norepinephrine: nor; dopamine: dop; epinephrine: epi; rosmarinic acid: RA; rosmarinic acid analog with an amide link, RA-amide; epigallocatechin gallate, EGCG. Statistics were conducted using one-way ANOVA, multiple comparisons test; *p <0.05. (E,F) ThT and TEM assays confirm that the anti-amylin amyloid activities of norepinephrine require aerobic exposure. Notice lack of fibril presence under aerobic condition with norepinephrine treatment.

Multiple small molecule amyloid inhibitors, many of which are catechol-containing polyphenols, perturb or “remodel” unaggregated and/or pre-aggregated amyloid species into denaturant-resistant aggregates that displayed broad-range molecular weights; characterized as “smear-type” distributions on SDS-PAGE gels (Ehrnhoefer et al, 2008; Hong et al, 2008; Bieschke et al, 2010; Palhano et al, 2013; Wu et al, 2017). Using RA and norepinephrine as two representative cases, treatment with a reducing reagent such as cysteine nearly eliminated their amyloid remodeling activities in a dose dependent manner (Figs. 3A & 3B). Similar effect was observed with glutathione and cystamine as well, and importantly, with other catechol-containing compounds including EGCG and dopamine, but not negative controls phenol and a non-catechol amyloid inhibitor, curcumin (Fig. 3C).

**Figure 3.**
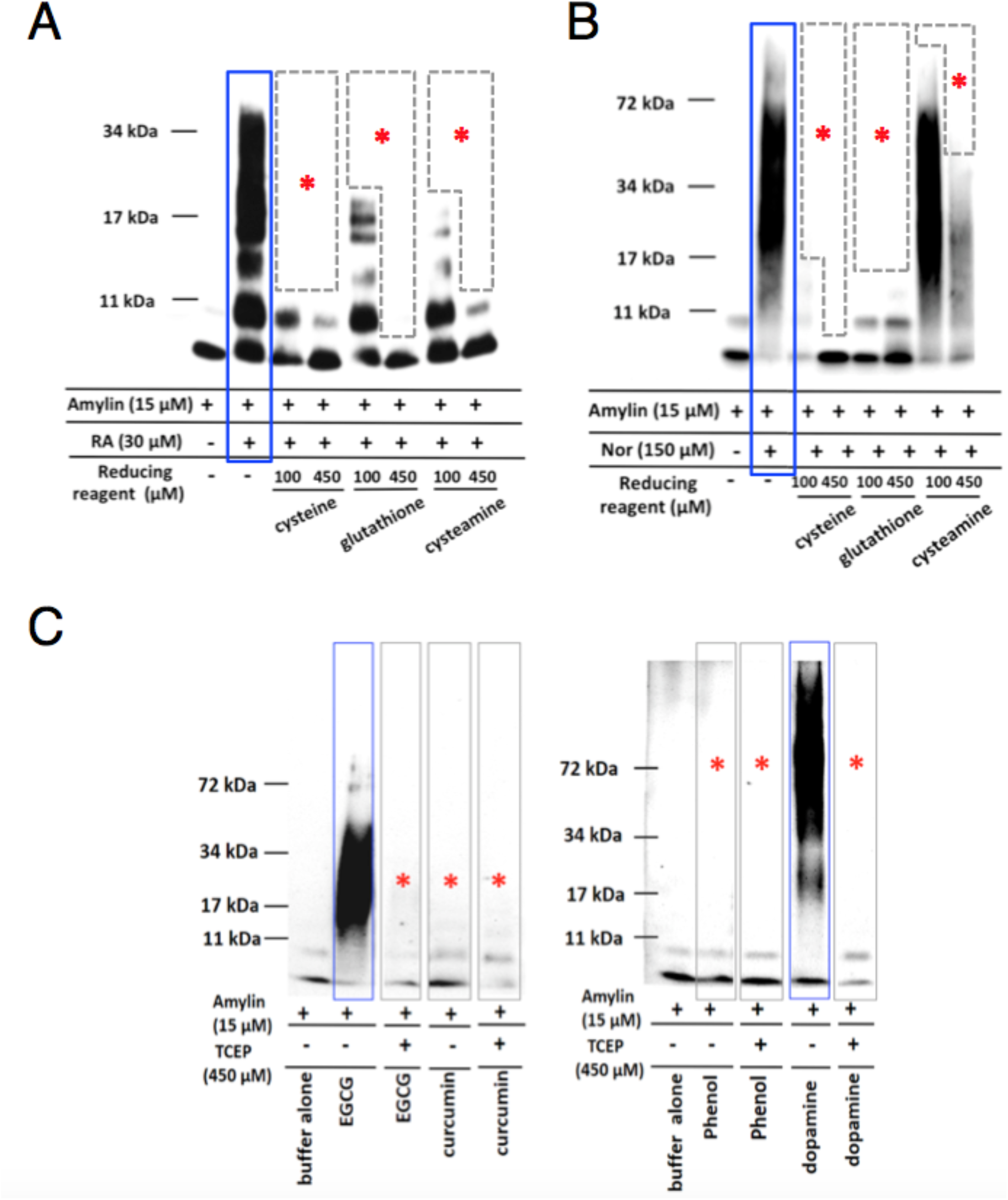
Blockage of amyloid remodeling activities of selective catechol-containing inhibitors by reducing agents. (A) Dose response effects of reducing agents cysteine, glutathione, and cysteamine on inhibiting RA (panel A) and norepinephrine (panel B) amyloid remodeling activities. Two different doses, 100 μM and 450 μM of reducing agents were used. (C) Control experiments demonstrate that reducing agent TCEP completely prevented the remodeling activities of amylin amyloid by catechol-containing inhibitors epigallocatechin gallate (EGCG) and dopamine but by neither a negative control, phenol, nor non-catechol inhibitor, curcumin. Amylin concentration used was 15 μM. The concentrations of reducing agent to inhibitors or control were 10:1 molar ratio (i.e., 450 μM TCEP, 45 μM of phenol, EGCG, or dopamine). In all panels, positive remodeling activities are highlighted in blue boxes and negative or mitigated remodeling activities are indicated by red asterisks.

The chemical changes that occur during autoxidation were investigated by both UV-Vis and liquid chromatography-mass spectrometry (LC-MS) approaches. UV-Vis time course spectra confirmed that chemical changes were only detectable under aerobic conditions, and occurred coincidently with their enhanced anti-amyloid activities (Figs. 4A, S4A, and S4B; highlighted by asterisks). Such chemical changes were reflected in broad UV absorption spectra changes, particularly in in region of 300-350 nm (Fig. 4A and Fig. S4). In an effort to identify the oxidized chemical species that contribute to the enhanced amyloid inhibition, we performed LC-MS analyses of aged norepinephrine. New species/peaks with increasing ion intensities over time (elution was collected at 0 - 96 hours) were detected in LC at the elution time between 0.85 min – 1.0 min (Fig. 4B). High-resolution MS identified four main ions within the norepinephrine sample (m/z peaks at 162.0548, 164.0715, 182.0458, and 301.1037 in Fig. 4C). While the known oxidized product of norepinephrine, noradrenochrome (C_8_H_5_NO_3_, 164.0342, [M+H]^+^) was anticipated product (Jimenez et al, 1984; Manini et al, 2007), the mass difference of 0.0373 from the observed 164.0715 does not support its presence. Thus all species remain unidentified. These ions may result from the acidic conditions of the LC separation as well as in-source reactions that may occur during the ionization process. Hence the features observed by LC-MS are presently considered signatures of the oxidation process and not the inhibitory species present in solution at neutral buffered pH. Nonetheless, these ions were present in the time course of autoxidation that correlate with norepinephrine anti-amyloid activities as exhibited by both ThT and TEM assays (Figs. 2E & 2F). These data agree well with studies that showed catecholamine oxidation products were effective anti-amyloidogenic agents against α-synuclein (Li et al, 2004) and tau (Soeda et al, 2015). Collectively, our data demonstrate that autoxidation is a general pathway enhancing the anti-amyloid activities of catechol-containing compounds (Fig. 4D), as we proposed in our specific investigation on baicalein (Velander et al, 2016). Redox states are therefore key factors that modulate the activities of a large number of catechol-containing amyloid inhibitors.

**Figure 4.**
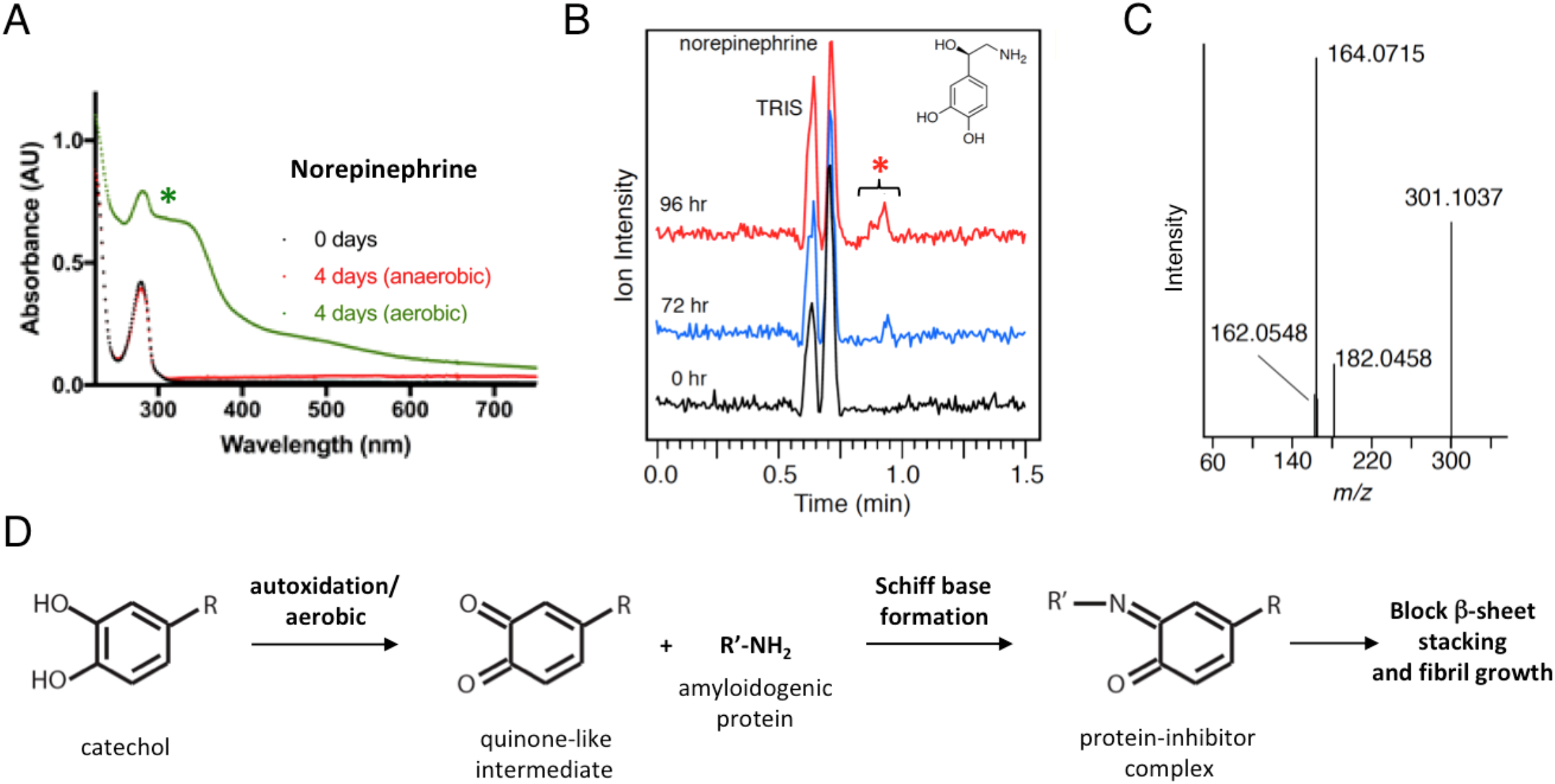
UV-Vis and LC-MS characterization of oxidized products of norepinephrine and model for amyloid inhibition by autoxidized catechol compounds. (A) Significant oxidative changes incurred for norepinephrine with 4 days exposure to aerobic condition, but not in anaerobic condition, in UV-Vis absorption spectra. (B,C) Freshly dissolved (time 0) or aged Norepinephrine samples (72 or 96 hours) were characterized by LC-MS. Visual inspection of the chromatograms showed a new peak at 0.85 – 1.0 minutes in the 72 and 96 hour sample that was not present at 0 hours. MS (positive mode) of this fraction identifies three species that are likely norepinephrine oxidized byproducts that include [M+ H^+^] 164.0715, 182.0458 and 182.0458. Albeit amylin peptide was absent in the LC-MS, similarly aged norepinephrine samples also show activity in TEM, ThT and gel remodeling assays (Figures 2 and 3). Such activity was not associated with non-aged norepinephrine, which importantly did not undergo oxidative chemical change. The latter demonstrates that at minimum, these species are likely associated with and may contribute to norepinephrine anti amyloid activities. Statistics were conducted using one-way ANOVA, Sidak’s multiple comparisons test; *p <0.05, ** p <0.01. (D) A general model of how autoxidation or aerobic conditions enhance the activities of catechol-containing protein amyloid inhibitors, as we originally proposed in our specific investigation on baicalein (Velander et al, 2016).

A variety of covalent mechanisms can readily explain the observed effects of autoxidation for catechol-mediated remodeling and the corresponding strong stability of the observed remodeled aggregates. Free radical cycling occurring during catechol autoxidation could directly cycle through nearby interacting amyloid proteins that subsequently lead to protein-protein and/or protein-compound adducts (Meng et al, 2009). Alternatively, remodeled aggregates may also form through covalent interactions between electrophilic o-quinone oxidized byproducts of catechol parent compounds and amyloid protein side chain amines (Zhu et al, 2004; Li et al, 2004; Hong et al, 2008; Meng et al, 2009; Popovych et al, 2012; Sato et al, 2013; Velander et al, 2016). Regardless of the exact mechanism, these data are consistent with the hypothesis that the presence of oxygen facilitates autoxidation, which in turn enables catechol-containing compounds to engage in amyloid remodeling activities that leads to what are presumably innocuous off-pathway, non-toxic aggregates (Ehrnhoefer et al, 2008; Hong et al, 2008; Bieschke et al, 2010; Wu et al, 2017). It remains to be determined whether amyloid remodeling is merely phenomenological in nature or if it represents a key mechanism essential for inhibitor-mediated anti-amyloid activities. Built from previous work (Zhu et al, 2004; Li et al, 2004; Pickhardt et al, 2005; Hong et al, 2008; Meng et al, 2009; Velander et al, 2016; Joshi et al, 2016; Velander et al, 2017; Wu et al, 2017), our data suggest that (i) catechol-containing compounds as well as redox related anthraquinones represent a broad class of amyloid inhibitors capable of targeting amyloid from multiple sequence-unrelated amyloidogenic proteins, and (ii) the redox states of catechol-containing compounds are a key determinant of their anti-amyloid activities.

We and several other research groups have shown that catechol groups in the catechol-containing polyphenols and flavonoids play key roles in amyloid inhibition (Caruana et al, 2011; Sato et al, 2013; Soeda et al, 2015; Velander et al, 2016; Wu et al, 2017). Extended from individual compounds, we have shown here that catechols and redox-related quinones/anthraquinones are a broad class of amyloid inhibitors. To our knowledge, identification of a class of compounds using systematic screens against multiple prototype amyloidogenic proteins has yet to be reported. The catechol moiety is a common structural component of many natural products, broadly classified as polyphenols, many of which have anti-amyloid and anti-aging activities (Cao & Raleigh, 2012; Ono et al, 2012; Yamada et al, 2015; Velander et al, 2017). Our results thus provide mechanistic insight into why this class of large number of natural compounds, often enriched in healthy diet, is active in neutralizing toxic amyloids, suggesting a combination of redox and structural modeling is a viable path to new anti-amyloid structures with enhanced activities. Catechol-containing natural products offer the advantage of low cost and potentially low side effects, inasmuch offering significant potential as anti-aging nutraceuticals.

Catechol and related quinone motifs are classified by Baell et al. as Pan Assay INterference compoundS (or PAINS) in drug discovery (Baell & Holloway, 2010; Baell, 2016; Baell & Nissink, 2018). However, validity of such overly broad classification is debatable as the authors conceded that about 5% of FDA-approved drugs contain PAINS-recognized substructures (Baell & Nissink, 2018). For an example, EGCG, a well-established amyloid inhibitor (Ehrnhoefer et al, 2008; Bieschke et al, 2010; Palhano et al, 2013) that has undergone numerous clinical trials, falls into the PAINS category, because it contains the catechol motif. Nevertheless, caution has been exercised in our work to validate with multiple orthogonal *in vitro* and cell-based assays. ThT fluorescence assays for example, one of the most commonly used assays for screening amyloid inhibitors, could lead to false positives with certain hydroxyflavones (Noor et al, 2012). Orthogonal secondary assays, such as TEM analysis, PICUP, and cell-based assays, allowed us to follow up on the hits generated from ThT fluorescence-based primary screenings, minimizing such false positives.

Multiple structural studies suggest that there are different conformers and/or molecular polymorphism in multiple amyloidogenic proteins including amylin, Aβ, tau, α-synuclein, and TDP-43 (Wiltzius et al, 2009; Tycko, 2015; Qiang et al, 2017; Guenther et al, 2018; Seidler et al, 2018; Cao et al, 2019; Arakhamia et al, 2020). Moreover, many studies indicate that unique amyloid conformers may faithfully propagate morphologically and biochemically between cells and tissues, in a prion-like manner (Frost et al, 2009; Watts et al, 2014; Strang et al, 2018). These experimental observations suggest that the accumulation and assembly of various protein aggregates in many protein misfolding diseases is not totally driven by the amino acid sequences of aggregation-prone molecules; instead, they may be governed by the precise cellular and pathological environment of aggregation conditions. It will be interesting to identify different conformers, key environmental factors, protein modifications, and correlate them with their cytotoxicity. Such structure and function classification will greatly facilitate the identification of specific pharmacophores for structure-based inhibitor design in the future.

## Materials and Methods

### Peptides and Chemicals

Synthetic amidated human amylin was purchased from AnaSpec Inc. (Fremont, CA) and the peptide quality was further validated by the Virginia Tech Mass Spectrometry Incubator. Hexafluroisopropanol (HFIP) and thioflavin T (ThT) were purchased from Sigma Aldrich (St. Louis, MO). Small molecule amyloid inhibitors and relevant control compounds were purchased from Toronto Research Chemicals (North York, ON, Canada), Fisher Scientific Inc. (Hampton, NH), Sigma-Aldrich Corp. (St. Louis, MO) or Cayman Chemical Company (Ann Arbor, MI). Dulbecco’s phosphate buffer saline (DBPS) pH 7.4, was purchased from Lonza (Walkersville, MD). Black 96-well non-stick-clear-bottom plates and optically clear sealing film were purchased from Greiner Bio-one (Germany) and Hampton Research (Aliso Viejo, CA) respectively. 300 mesh formvar-carbon-coated copper grids and uranyl acetate replacement solution (UAR) were purchased from Electron Microscopy Sciences (Hatfeild, PA).

Lyophilized amylin powder (0.5 mg) was initially dissolved in 100% HFIP at a final concentration of 1-2 mM. The additional lyophilizing step was employed to eliminate traces of organic solvents, which have been shown to affect amylin aggregation. Aliquots were either lyophilized again prior to use in cell-based assays or dissolved directly into DPBS, 10 mM phosphate buffer pH 7.4 or 20 mM Tris-HCl pH 7.4 for all amylin amyloid-related *in vitro* assays. All remaining 1-2 mM stocks in 100% DMSO were stored at −80 °C until later use. The lyophilized powder from all compounds and ThT were dissolved in DMSO (10 mM) and distilled water or relevant buffer (1-4 mM). These stocks were stored at −20 °C until later use. Residual DMSO in the final samples used for all *in vitro* assays ranged from 0-9.5%. We determined that these DMSO concentrations had negligible effects on amylin amyloid aggregation as reflected by ThT fluorescence, TEM, PICUP assay, and inhibitor-induced amylin amyloid remodeling assays.

UV-Vis absorption spectra were collected on a Varian Cary 50 UV-Vis spectrophotometer. Equal concentration of catechol, norepinephrine, and rosmarinic acid was used for each compound. Aerobic and anaerobic conditions referred in each case as exposure to air or incubation in an anaerobic chamber (described below) at the specified duration.

### Thioflavin-T Fluorescence Screening Assays

Fluorescence experiments were performed using a SpectraMax M5 plate reader (Molecular Devices, Sunnyvale, CA) as described in our previous works (Velander et al, 2016; Wu et al, 2017). All kinetic reads were taken in non-binding all black clear bottom Greiner plates covered with optically clear sheets to minimize the evaporation of the solution in each well and stirred for 10 seconds prior to each reading. All experiments were repeated three times using peptide stock solutions from the same lot.

#### (a) Screening against amylin amyloid

Amylin stock preparation was initiated by dissolving lyophilized human amylin powder in 100% HFIP. After at least 2 hours of incubation at room temperature, HFIP was removed and the resulting amylin powder was re-dissolved in 100% DMSO and quantified by a standard BCA micro plate assay (i.e. HFIP is extremely volatile and difficult to accurately pipette; thus, DMSO is a better solvent for peptide transfer. Initial HFIP dissolution was employed because it has been suggested to aid in removal of trace contaminants that may be present within lyophilized batches of peptide). Solutions of 400 μM amylin (100% DMSO) were subsequently diluted with ThT in 1X DPBS and aliquoted to reading plate wells containing 0.45 μL of compound or vehicle buffer controls to a final volume of 15 μL. Final concentrations prior to amylin amyloid aggregation were 10 μM amylin, 30 μM compound and 30 μM ThT. Aggregation reactions were incubated at 25°C until the approximated plateau fluorescent readings. All plates were continuously monitored for ThT fluorescence every 10 minutes (plates were shaken briefly prior to each read) at excitation 444 nm and emission 490 nm until the plateau phase was reached.

#### (b) Screening against 2N4R tau

Purified recombinant 2N4R tau (the largest and the most studied human tau isoform) initially dissolved in 100% 1X PNE buffer, was buffer exchanged with 1X HEPES buffer (10 mM HEPES, 30 mM NaCl, pH 7.4). Amyloid aggregation was initiated in the presence or absence of compound by diluting tau protein into the wells of the reading plates to a final 34.4 μM of tau, 12.5 μM ThT and 0.06 mg/ml of heparin (served to promote tau amyloid formation) in each well. For compound treated wells, final molar concentration of compound to tau was 2:1. Tau aggregation reactions were incubated at 37°C and the fluorescent readings were taken at excitation 444 nm/emission 490 nm.

#### (c) Screening against Aβ42 peptide

In contrast to the ThT fluorescence assays for amylin and 2N4R tau, non-continuous, two-point measurements were taken to represent starting and ending (at plateau) fluorescent signals for Aβ42. Unaggregated solutions of Aβ42 were typically prepared by dissolving lyophilized Aβ42 powder with 100% HFIP. Subsequent peptide quantification of this solution was estimated based on a standard micro plate BCA assay. Initial stock solution preparation of Aβ42 for the ThT assay were prepared by evaporating HFIP treated stocks followed by a modified two-step aqueous dissolution process as previously described: Herein, HFIP evaporated stocks of Aβ42 were dissolved in 60 mM NaOH. Next, these solutions were further diluted in 1X DPBS such that the final solution consisted of approximately 2.4 mM NaOH (i.e. 4% 60 mM NaOH, 96% 1X DPBS). Immediately after the addition of 1X DPBS, 6.48 μL aliquots of Aβ42 and 0.52 μL aliquots of compound or buffer treated controls were distributed to the ThT reading plates. All plates were sealed to prevent evaporation and allowed to incubate at 37°C until the estimated plateau of aggregation (24 hours). Next, the reading plates were centrifuged prior to receiving 14 μL of a 45 μM ThT solution prepared in 50 mM Glycine-NaOH, 8.6 pH. Finally, all plates were read at excitation 444 nm/emission 490 in order to estimate plateau phase amyloid aggregation as indicated by ThT fluorescence. Note, the final concentration of Aβ42 prior to being titrated with ThT, was approximately 12 μM, at a 3:1 molar ratio of compound to Aβ42.

The NIH Clinical Collection (NIHCC) library contains a total of 700 FDA-approved drugs and investigational compounds with diverse chemical structures. These small molecules have a history of use in human clinical trials. The collection was assembled through the NIH Molecular Libraries and Imaging Initiative. The library was supplied in 96-well plates in DMSO. For screening, 384-well plate was used. Bravo liquid handler with 96-channel disposable tip head (Agilent Technologies, Wilmington, DE) was used to aid sample transfer.

Total volume in each well was 15 μl. Residual DMSO (<5.4%) in each well had negligible effect on the fluorescent signals.

### Aggregation Assays in Anaerobic Chamber

All reagents including buffers, protein and compounds were made anaerobic by flushing with nitrogen prior to being placed within an anaerobic chamber (glove box). H_2_ gas mixed with inert gas is circulated through metal catalyst to remove O_2_ gas in the chamber. Oxygen level was maintained between 15-50 ppm within the anaerobic chamber. Typical aggregating conditions within the chamber were maintained at room temperature (25-30 °C). All anaerobic chamber-based ThT fluorescence assays were discontinuous, two point measurements (starting point, ending or plateau point).

### ThT Fluorescence-Based Amyloid Remodeling Assay

Similarly with regular ThT fluorescence amyloid formation assays, remodeling assays extended the monitoring of the fluorescence signals continuously after an inhibitor or a control compound was spiked into the aggregation system (amyloidogenic protein, ThT, buffer, heparin the case of tau protein). The volume of spiked compound was made to 5% of total sample volume in each well such that the baseline fluorescence signal was minimally changed.

### Transmission Electron Microscopy (TEM) Analysis

TEM images were obtained by a JEOL 1400 microscope operating at 120 kV. Samples consisting of 30 μM amylin (20 mM Tris-HCl, 2% DMSO, pH 7.4) in the presence of drug or vehicle control were incubated for ≥48 hours at 37 °C with agitation. Prior to imaging, 2-5 μL of sample were blotted on a 200 mesh formvar-carbon coated grid for 5 minutes and then stained with uranyl acetate (1%). Both sample and stain solutions were wicked dry (sample dried before addition of stain) by filter paper. Qualitative assessments of the amount of fibrils or oligomers observed were made by taking representative images following a careful survey of each grid (>15-20 locations on each grid were surveyed).

### Gel-Based Amyloid Remodeling Assay

Vehicle control or specified compounds were spiked into freshly dissolved amylin samples (containing amylin and buffer). Thereafter amylin aggregation was allowed to proceed for 3 days. Final amylin concentration was 15 μM that included 45 μM of compound (drug:amylin molar ratio was 3:1). After 3 days, these samples were vacuum dried and re-dissolved in 6.5 M urea containing 15 mM Tris and 1X SDS Laemmeli sample buffer, boiled at 95 °C for 5-10 minutes and subjected to SDS-PAGE followed by Western blot analysis with anti-amylin primary antibody (T-4157, 1:5000, Peninsula Laboratories, San Carlos, CA). All gel-based amyloid remodeling assays were repeated at least twice.

### Photo-Induced Crosslinking of Unmodified Proteins (PICUP) Assay

Amylin aliquots from a master mix in 10 mM phosphate buffer, pH 7.4 were added separately to 0.6 mL eppendorf tubes containing small molecule inhibitors or DMSO vehicle loaded controls. Crosslinking for each tube was subsequently initiated by adding tris(bipyridyl)Ru(II) complex (Ruby) and ammonium persulfate (APS) (Typical amylin:Rubpy:APS ratios were fixed at 1:2:20, respectively, at a final volume of 15-20 μL), followed by exposure to visible light, emitted from a 150-Watt incandescent light bulb, from a distance of 5 cm and for a duration of 5 seconds. The reaction was quenched by addition of 1X SDS sample buffer. PICUP results were visualized by SDS-PAGE (16% acrylamide gels containing 6 M urea), followed by silver staining. Final concentrations for all PICUP reactions included 30 μM of amylin and 150 μM of each compound.

### Liquid Chromatography – Mass Spectrometry (LC-MS) Analysis

Analyses were performed on a Waters Synapt G2-S HDMS interfaced with an Acquity I-Class UPLC system. Freshly dissolved (time 0) or aged (exposure to air for 72 or 96 hours) norepinephrine samples at a concentration of 37.5 μM (diluted in a mixed solvent of 2:98, acetonitrile:water, v/v) were characterized by positive mode LC-MS for its oxidative products. Norepinephrine was reliably detected on the reverse phase column with a retention time of 0.7 minutes. Visual inspection of the chromatograms showed a new peak at retention time between 0.85-0.95 minutes in the 72 and 96 hour sample that was not present at 0 hours. Oxidized species (new peaks) were characterized by MS/MS analysis.

The binary solvent system was composed of 0.1% formic acid in water (A) and 0.1% formic acid in acetonitrile (B). A 10-minute gradient was used for the analysis with the following conditions: initial and hold for 30s at 1%B, a linear gradient to 90%B at 8 min, hold at 90%B to 8.5 min and return to initial condition at 9 min. Sample volumes for MS analysis were between 2-4 μl and injected onto a Waters UPLC BEH C18 (1.7 μm, 2.1 mm x 50 mm) held at 35 °C for MS analysis. Source conditions for the mass spectrometer were: capillary 2.8 kV, source temperature 125 °C, sample cone 30V, source offset 80V, desolvation gas 500 L/Hr, desolvation temperature 400 °C, cone gas 50 L/Hr, and nebulizer gas 6 par. The *m/z* scan range was 50-1800 for MS analysis and 50-600 for MS/MS analysis and a collision energy ramp of 15-40 eV was used for MS/MS analysis.

### Statistical Analysis

All data are presented as the mean ± S.E.M and the differences were analyzed with a one-way analysis of variance followed by Holm-Sidak’s multiple comparisons (amylin kinetics) or unpaired Student’s *t* test. These tests were implemented within GraphPad Prism software (version 6.0). *p* values < 0.05 were considered significant.

## Acknowledgments

This work was supported in part by the Hatch Program of the National Institute of Food and Agriculture, USDA (BX and RFH), Commonwealth Health Research Board Grant 208-01-16 (BX and SZ), Diabetes Action Research and Education Foundation Grant (BX), Awards No. 16-1, 18-2 and 18-4 from the Commonwealth of Virginia’s Alzheimer’s and Related Diseases Research Award Fund, administered by the Virginia Center on Aging, School of Allied Health Professions, Virginia Commonwealth University (BX, LW, and SZ), NIH grants R03AG061531 (BX) and R01AG058673 (SZ), Alzheimer’s Association/Michael J. Fox Foundation grant BAND-19-614848 (BX), and Alzheimer’s Drug Discovery Foundation 20150601 (SZ). We thank Ms. Kathy Lowe at Virginia-Maryland Regional College of Veterinary Medicine for her excellent technical assistance in collecting TEM data. We thank Shradha Ladd for assistance in 2N4R tau recombinant protein purification. We thank Virginia Tech Center for Drug Discovery and Dr. Pablo Sobrado for instrument and NIHCC library access. We thank Prof. Jianyong Li for constructive discussions.

## Author Contributions

PV, LW, SBH performed the experiments, acquired and analyzed the data. NJV assisted with NIHCC library screening experiment and robotics usage. RFH designed LC-MS experiments and analyzed the data. BM, SZ contributed key reagents or facility and provided intellectual inputs in experimental design/method. BX conceived, organized, designed the experiments, and analyzed the data. BX, PV, LW, RFH wrote the paper.

## Additional Files

Supplementary Files

Three supplementary tables and four supplementary figures are attached.

## Supplemental Data

**Table S1.**
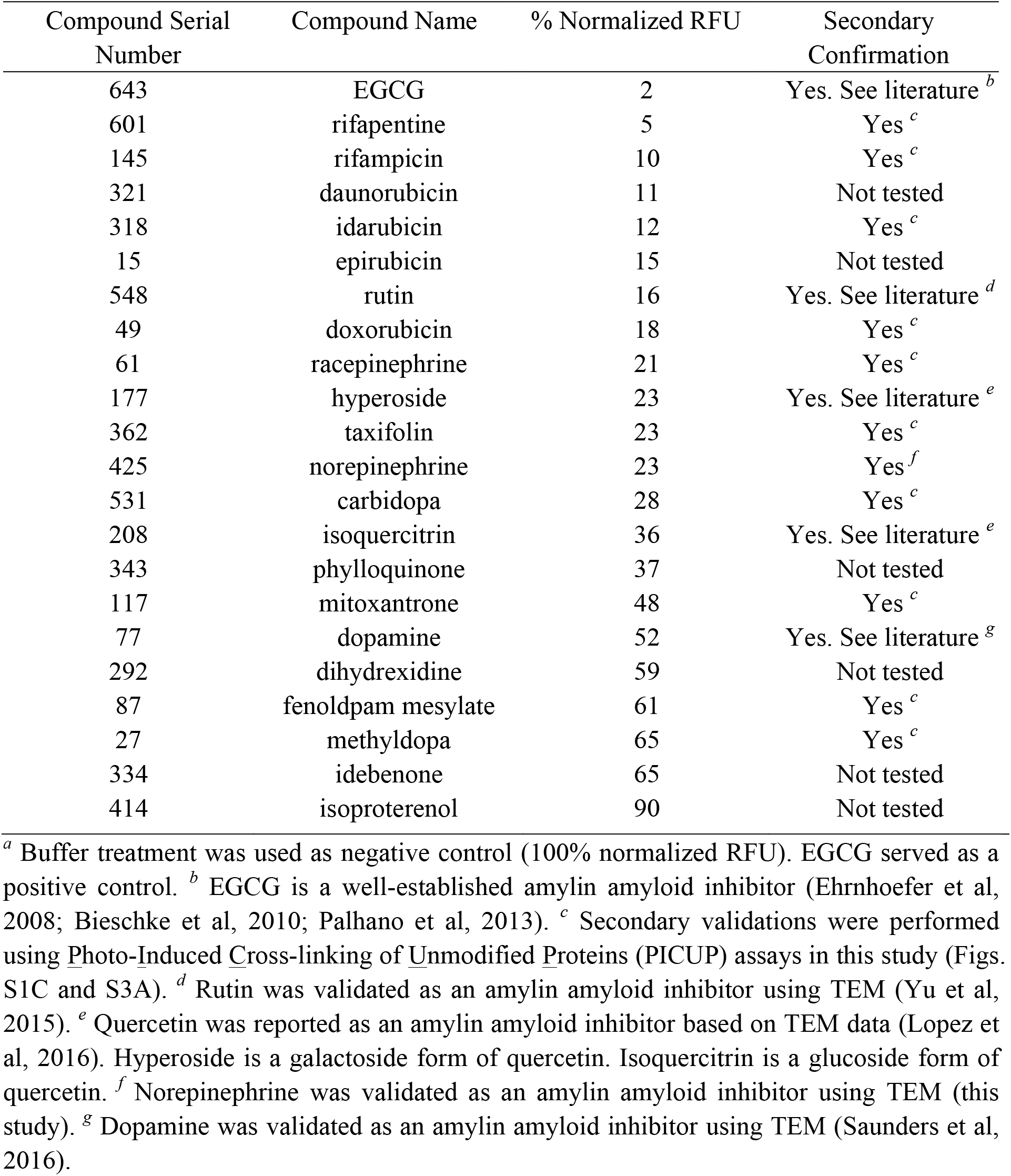
Catechol-containing compounds or redox-related anthraquinones from NIHCC library and their amylin amyloid inhibitory potencies in ThT fluorescent screen assays.^*a*^

**Table S2.**
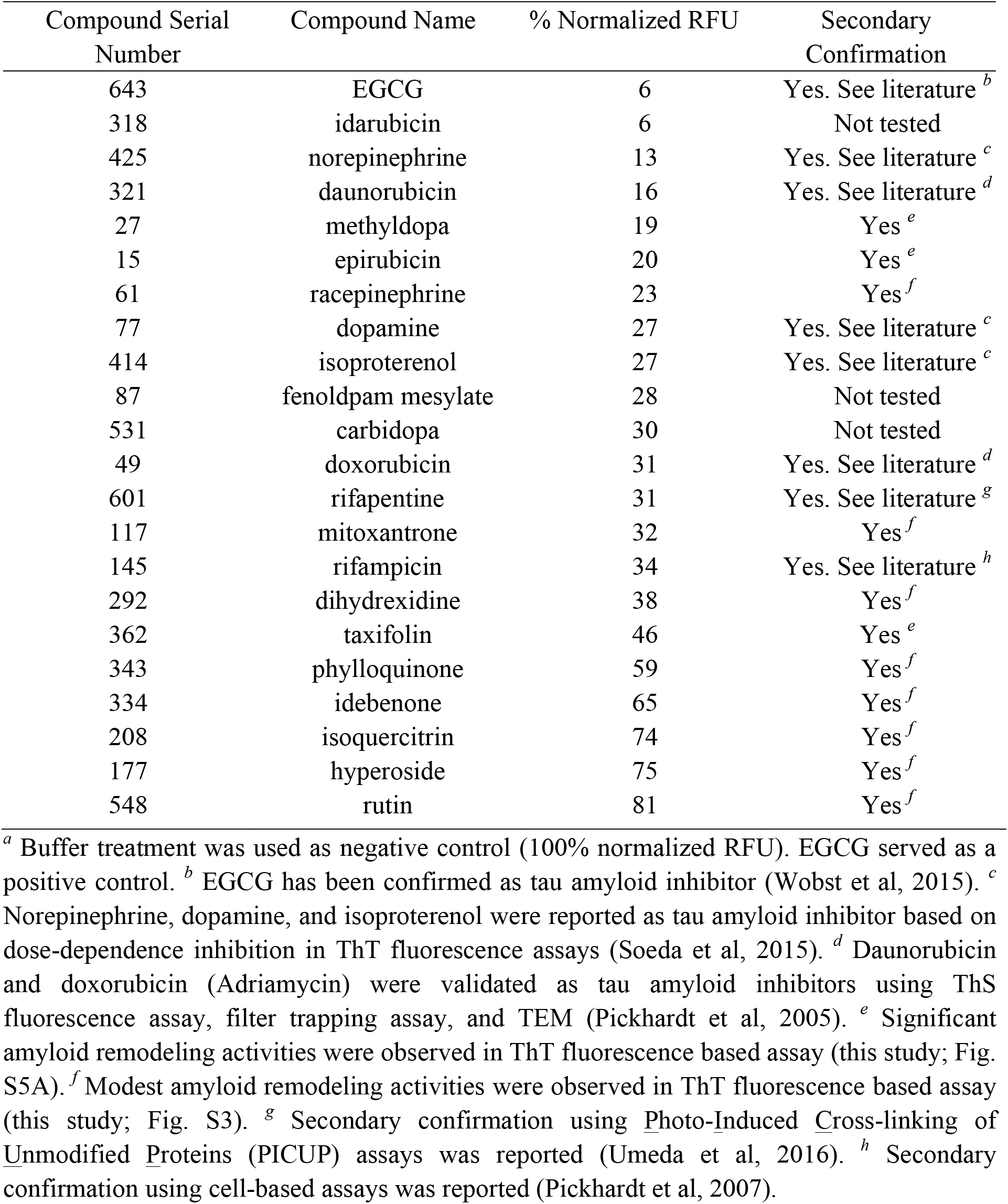
Catechol-containing compounds or redox-related anthraquinones from NIHCC library and their human tau (2N4R isoform) amyloid inhibitory potencies in ThT fluorescent screen assays.^*a*^

**Table S3.**
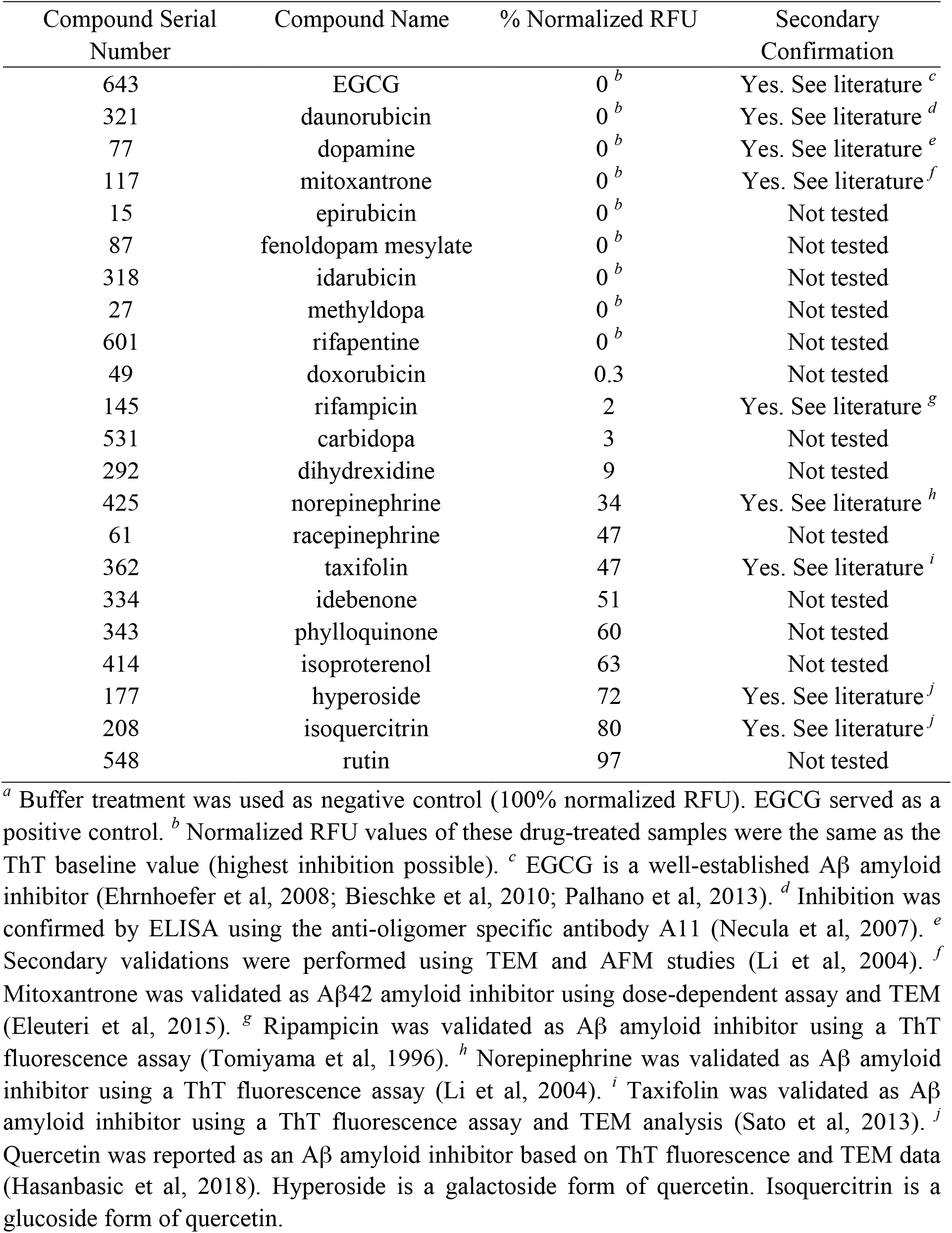
Catechol-containing compounds or redox-related anthraquinones from NIHCC library and their Aβ42 amyloid inhibitory potencies in ThT fluorescent screen assays.^*a*^

**Figure S1.**
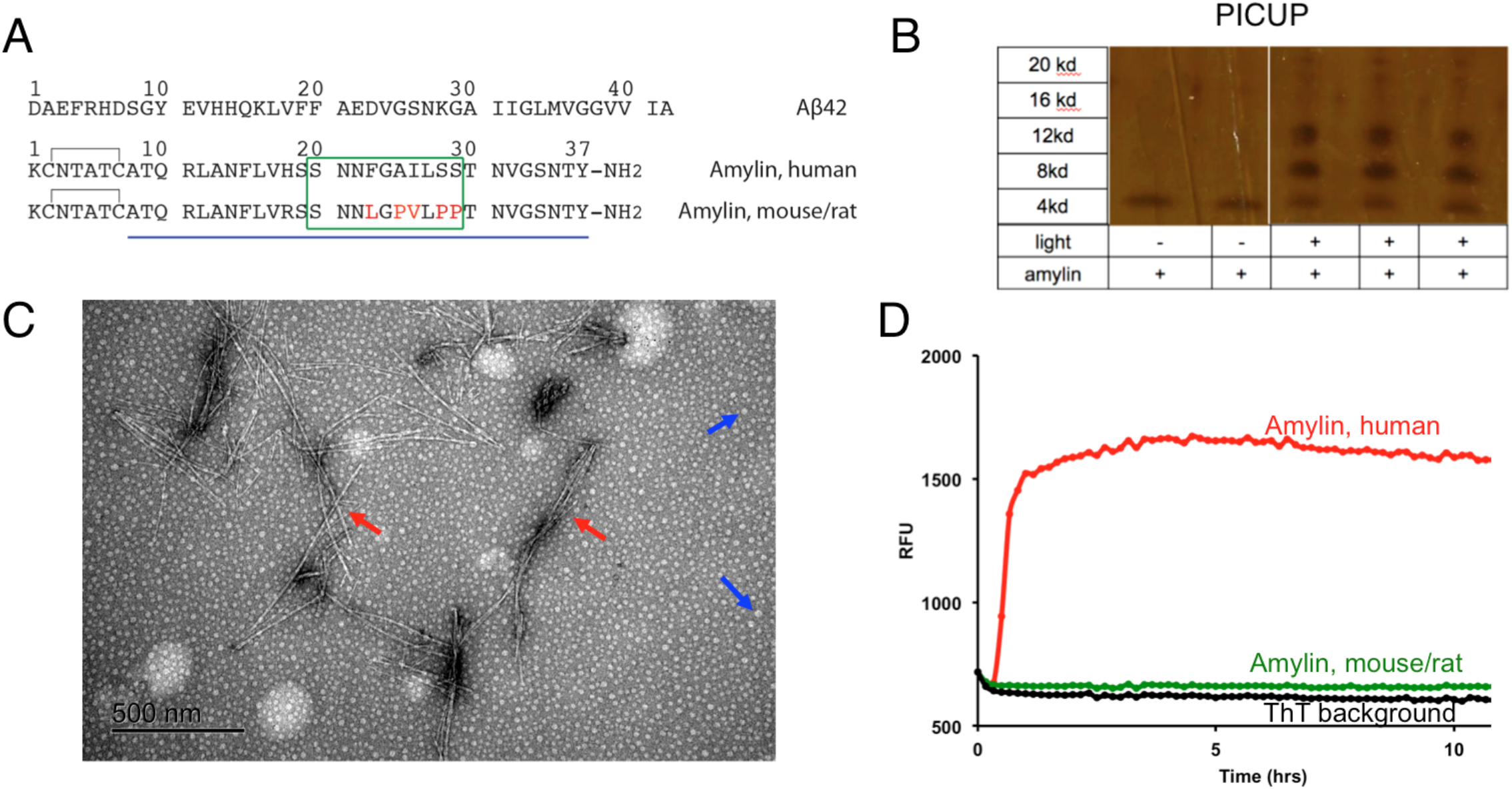
Amylin and Aβ42 sequences and biophysical assays for analyzing protein amyloids and oligomers. (A) Amino acid sequences of human and rodent amylin and amyloid β peptide (Aβ) are shown. The green box indicates a key region in amylin sequence (20-29) determining human amylin amyloid formation. Different amino acids between human and rodent amylin sequences are highlighted in red. (B) Proof-of-principle *in vitro* photo induced cross-linking of unmodified proteins (PICUP) assay that identifies stable amylin oligomers. The assay was analyzed with SDS-PAGE followed by silver staining. (C) Transmission electron micrographic (TEM) image of human amylin amyloid. Mature fibrils and spherical oligomers are indicated by red and blue arrows respectively. (D) ThT fluorescence assay showing human amylin amyloid formation, but not non-amyloidogenic rodent amylin. ThT only background is also shown.

**Figure S2.**
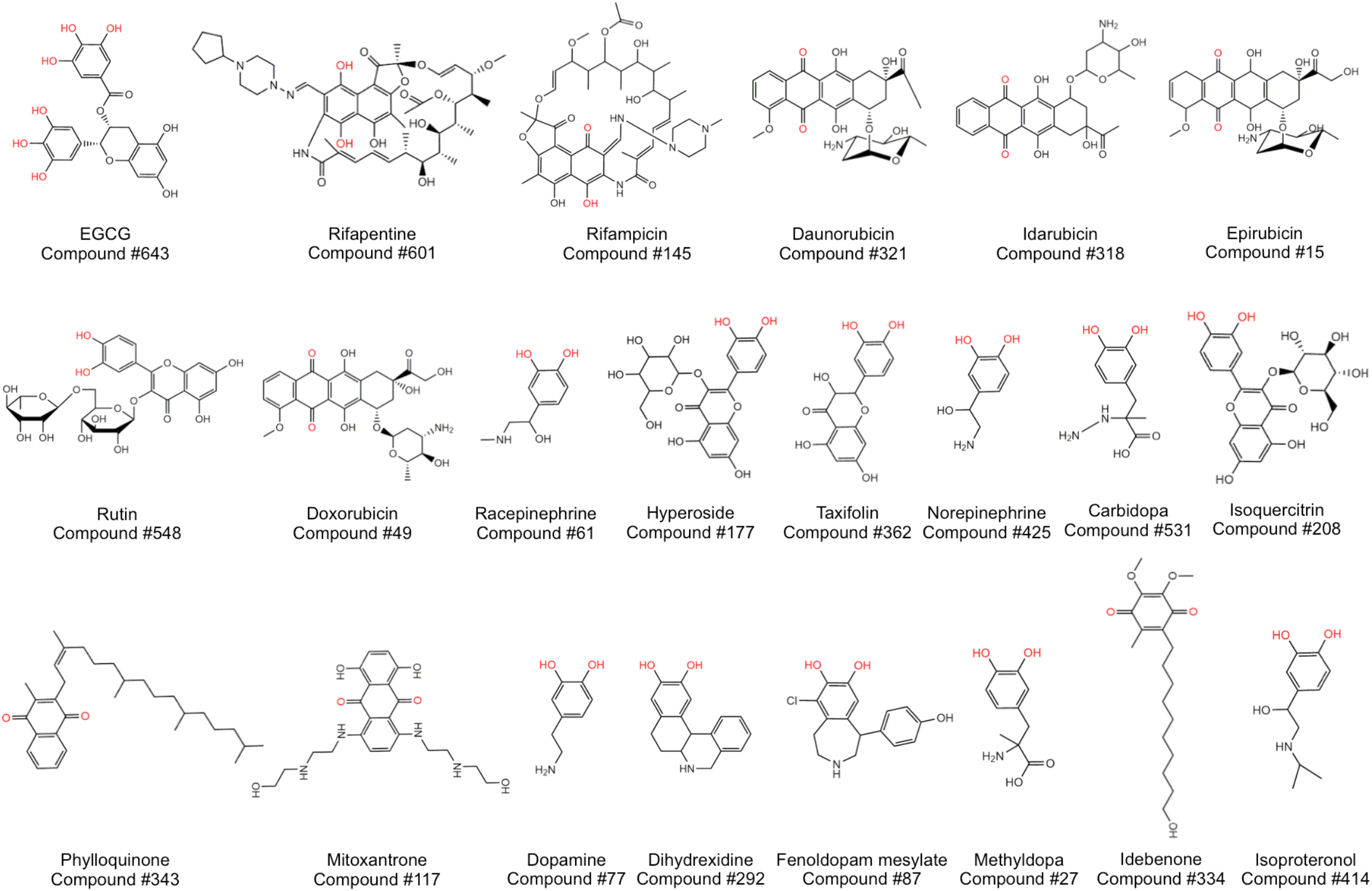
Chemical structures of catechols and redox-related quinones and anthraquinones identified from the NIHCC repurposing library of 700 drug and investigational compounds for their amyloid inhibitory activities of amylin, tau 2N4R and Aβ42. Relevant functional chemical groups are highlighted in red. Out of these 22 compounds, 21 compounds showed significant inhibitory effects against amylin amyloid (Table S1). Similarly, out of these 22 compounds 17 compounds have significant activities against tau 2N4R amyloid (Table S2) and 18 hits have significant activities against Aβ42 amyloid (Table S3).

**Figure S3.**
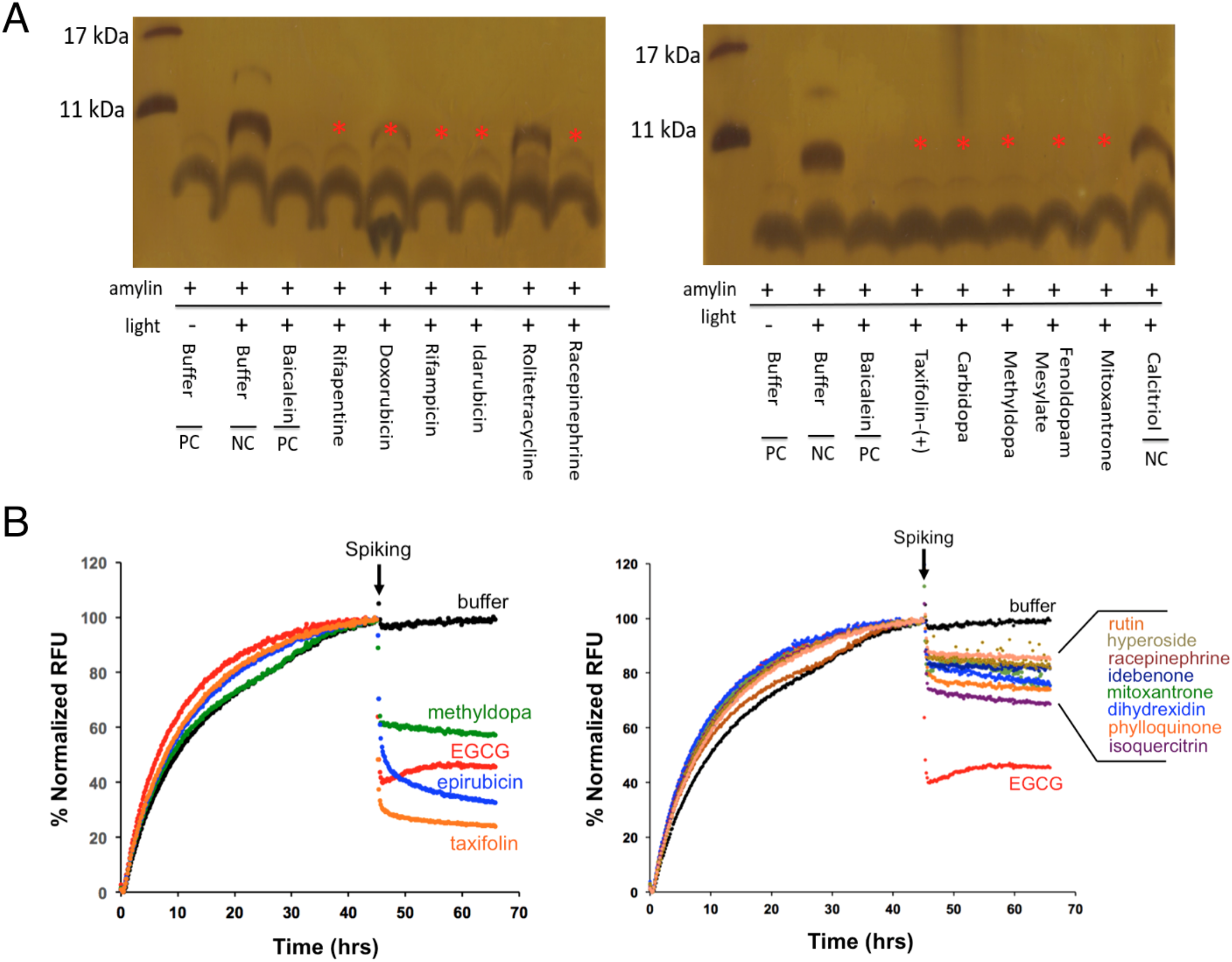
Secondary validation of selected top hits of catechols and quinones/anthraquinones from NIHCC library. (A) Secondary validation of amylin amyloid inhibitor hits by PICUP analysis. Amylin concentration was 30 μM and compound to amylin molar ratio was 5:1. Buffer and two drugs, rolitetracycline and calcitriol, were used as negative controls. Baicalein served as a positive control (Velander et al, 2016). Absence (or reduced intensity in the case of doxorubicin) of dimer formation bands (red asterisks) indicates their significant activities in blocking amylin oligomer formation. (B) Secondary validation of selected tau amyloid inhibitors by ThT fluorescence-based remodeling assay. Tau 2N4R isoform concentration was 30 μM and compound to tau 2N4R molar ratio was 2:1. Drugs are spiked at the indicated time (black arrows). Buffer (black lines) was used as a negative control. EGCG (red lines) served as a positive control. Compounds with strong remodeling effects were shown in the left panel and compounds with moderate remodeling effects were shown in the right panel.

**Figure S4.**
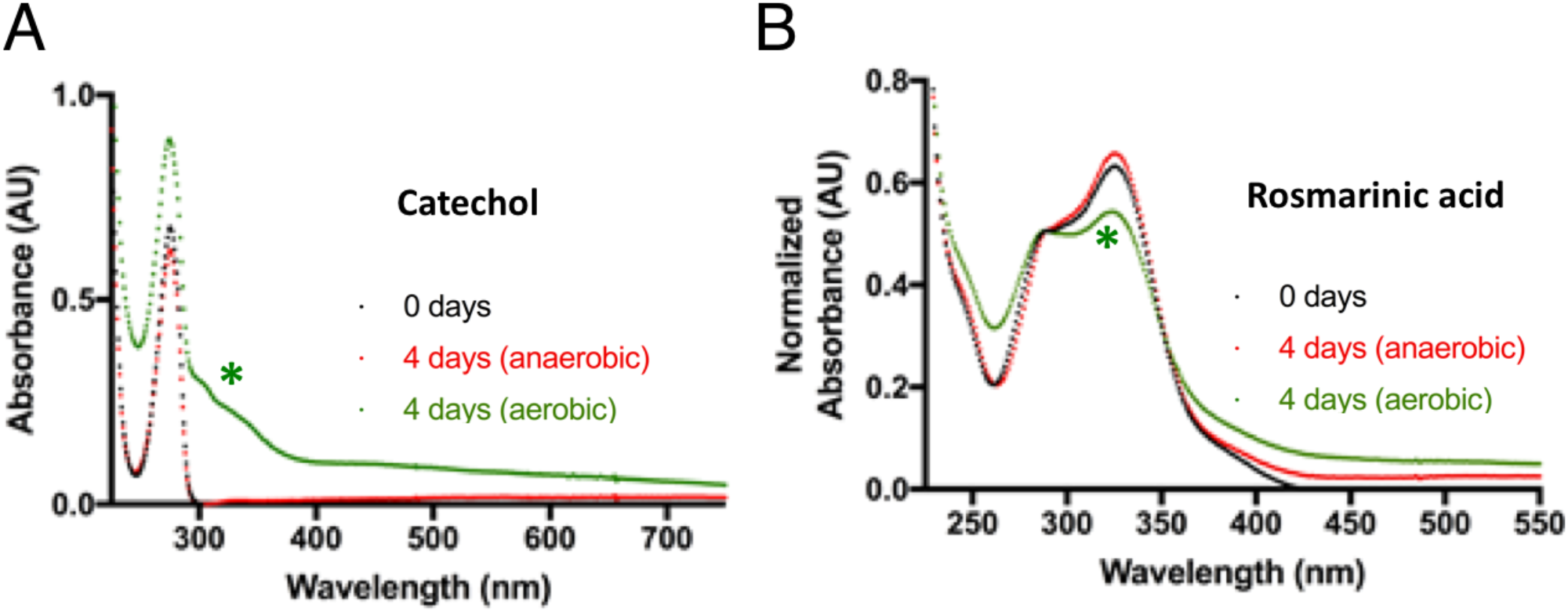
UV-Vis spectra reflect the oxidative changes incurred for aerobic exposed catechol or catechol-containing compounds, but not the same compounds under anaerobic condition. Significant spectra changes occurred for catechol (A) and rosmarinic acid (B) under aerobic condition (green traces) but not anaerobic conditions (red traces). The zero day (black trace) time point represents the compound spectra taken prior to undergoing incubation in either aerobic or anaerobic environments.

